# Should bionic limb control mimic the human body? Impact of control strategy on bionic hand skill learning

**DOI:** 10.1101/2023.02.07.525548

**Authors:** Hunter R. Schone, Malcolm Udeozor, Mae Moninghoff, Beth Rispoli, James Vandersea, Blair Lock, Levi Hargrove, Tamar R Makin, Chris I. Baker

## Abstract

A longstanding engineering ambition has been to design anthropomorphic bionic limbs: devices that look like and are controlled in the same way as the biological body (biomimetic). The untested assumption is that biomimetic motor control enhances device embodiment, learning, generalization, and automaticity. To test this, we compared biomimetic and non-biomimetic control strategies for able-bodied participants when learning to operate a wearable myoelectric bionic hand. We compared motor learning across days and behavioural tasks for two training groups: Biomimetic (mimicking the desired bionic hand gesture with biological hand) and Arbitrary control (mapping an unrelated biological hand gesture with the desired bionic gesture). For both trained groups, training improved bionic limb control, reduced cognitive reliance, and increased embodiment over the bionic hand. Biomimetic users had more intuitive and faster control early in training. Arbitrary users matched biomimetic performance later in training. Further, arbitrary users showed increased generalization to a novel control strategy. Collectively, our findings suggest that biomimetic and arbitrary control strategies provide different benefits. The optimal strategy is likely not strictly biomimetic, but rather a flexible strategy within the biomimetic to arbitrary spectrum, depending on the user, available training opportunities and user requirements.

## Introduction

In an iconic scene in science-fiction cinema, Luke Skywalker is shown examining his new bionic prosthetic hand (*1*). The device appears to, nearly perfectly, mimic a biological hand in its visual appearance and function. The control of the hand appears intuitive, such that Luke can immediately manipulate the individual digits with high dexterity. While today’s technology is far from Skywalker’s, the design of artificial prosthetic hands has steadily innovated appearance and functionality to be increasingly like a biological hand: from 16^th^ century iron-clad hands with no manipulability, 20^th^ century body-powered hook devices capable of simple grasping (*2*) to modern multi-gestural bionic hands that can be operated via EMG pattern-recognition control systems [Coapt LLC Complete Control system; Ottobock Myo Plus; for a review of available bionic prosthetic hands, see (*3*)]. Driving much of the previous research and development in prosthetics, as well as the future trajectory of this industry, is the long-standing engineering ambition to design anthropomorphic artificial limbs: devices that look like and are controlled in the same way as the biological body, i.e., *biomimetic* (*4*). Biomimetic design has also driven the development of more invasive human-machine interfaces such as artificial sensory feedback systems (*5*–*11*), as well as brain computer interfaces (*12*). Across these various approaches, biomimetic-inspired design in human-machine interfaces is predicated on the (largely untested) assumption that biomimetic devices potentially allow users to recruit pre-existing neural resources supporting the biological body to assist device control, thereby enhancing device learning, generalization, sense of embodiment, and automaticity. But are these assumptions that underlie biomimetic design valid?

If these assumptions are valid, we would expect the brain to integrate neural representations of external devices with the biological body to support an efficient recruitment of neural body resources. However, one growing body of evidence has suggested that this may not be feasible. Recent neuroimaging studies have found that individuals with extensive experience using a device as a hand replacement (prosthetic hands or expert grasping tools) neurally represent their devices *less* like a biological hand (i.e., more distinct representations), as compared to novices (*13*, *14*). So then, why should devices be designed to mimic the body if the brain doesn’t seem to process, even the most highly used, external devices in the same way as a biological body-part? Moreover, considering the stark differences between biological and modern bionic limbs (e.g., response time, dexterity, functionality, aesthetics, comfort/fit, weight, durability, sensory feedback), there are multiple ways in which biomimetic interfaces may actually introduce neurocognitive conflicts for users between pre-existing information/resources for the biological body and those for the artificial device (*15*). Lastly, considering that most surveys of prosthesis users report high rates of prosthesis dissatisfaction and complete device abandonment (*16*, *17*), a critical re-evaluation of the research priorities driving development of these devices is warranted. In particular, it is vital to evaluate non-biomimetic control strategies that prioritize other design considerations (such as user requirements) over explicit biomimetics.

Here, we compared biomimetic and non-biomimetic motor control strategies directly while participants learned to operate a bionic hand. As a striking alternative to biomimetic control, we incorporated an arbitrary (non-biomimetic) control strategy. Based on the neurocognitive assumptions underlying biomimetic design, this strategy should provide no direct benefits for the user. The primary rationale of the arbitrary strategy is to provide a complete contrast to biomimetic. To test bionic hand skill learning, we trained able-bodied participants (n=40) to use a wearable myoelectric bionic hand (Figure 1). We assessed motor learning on multiple bionic hand skills across 4 training days (2-3 hours per day) and 2 testing days for the training groups: Biomimetic (n=20; mimicking the desired bionic gesture with biological hand) and Arbitrary (n=20; mapping an unrelated hand gesture with the desired bionic gesture). After training, we assessed how well the learning for each control strategy would generalize to a novel control strategy. We also tested a control group (n=20) that received no bionic hand training (i.e., the Untrained group). Based on the assumptions underlying biomimetic-inspired design, one would predict that biomimetic control would provide an increased sense of embodiment, better performance, generalization, and more intuitive control. In contrast, due to potential neurocognitive conflict associated with biomimetic control, our (pre-registered) core prediction was that training using an arbitrary control strategy might show increased performance over training, as well as post-training generalization to a novel control mapping (for pre-registered predictions see https://osf.io/3m592/). In contrast, biomimetic control might provide specific advantages in short-term performance and automaticity (more intuitive control). Additionally, we predicted that, regardless of the control type, bionic hand skill learning would be associated with increased sense of embodiment and motor control over the course of training.

**Figure 1.**
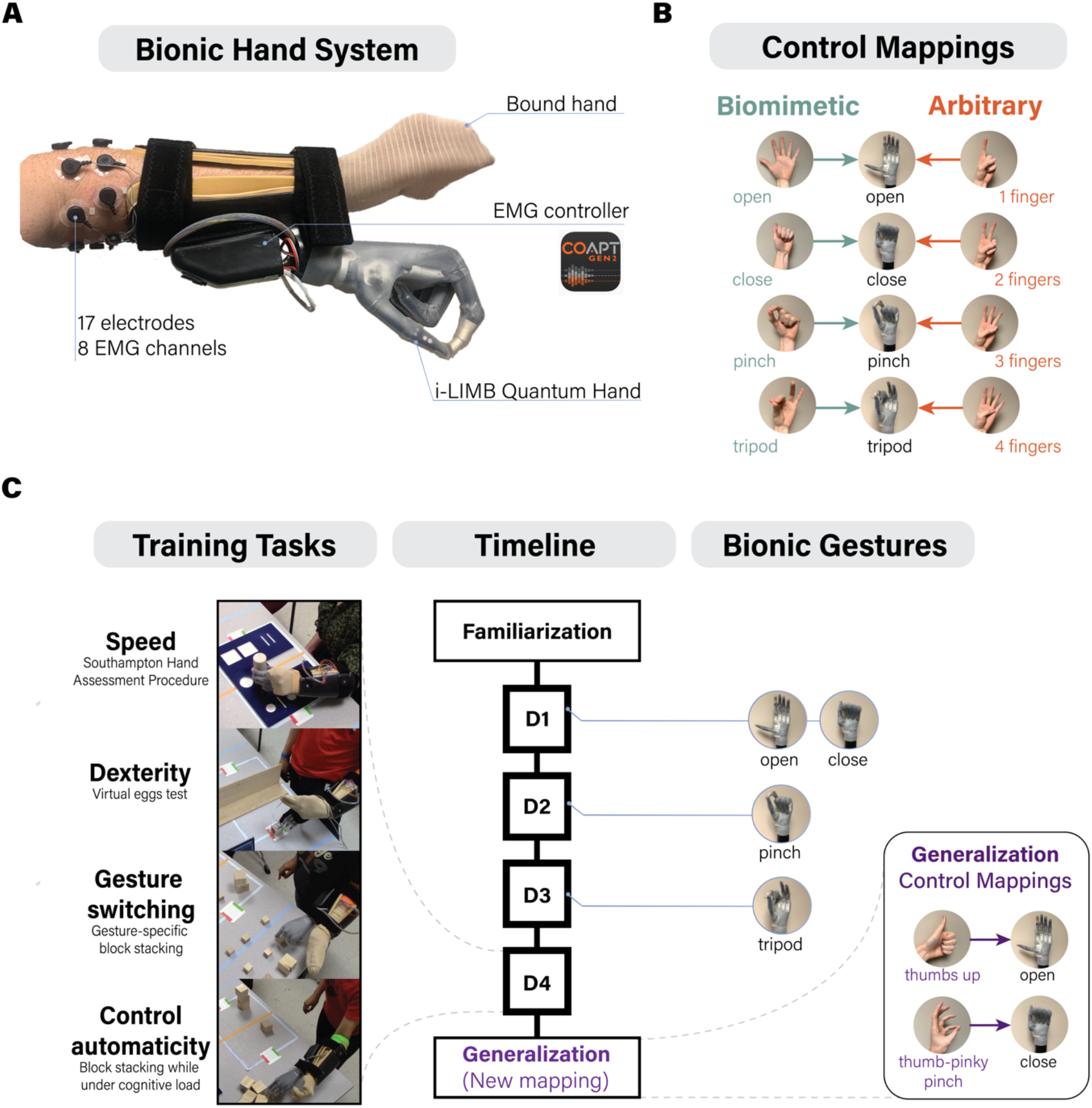
Experimental design of the study. **(A)** Bionic hand system attached to the participant’s left arm. The i-LIMB Quantum bionic hand is controlled by a Coapt pattern recognition controller (*18*) using signals from surface electromyography (EMG) electrodes (8 channels) positioned around the muscles of the forearm (for a detailed breakdown of device components, see Methods). The biological hand was bounded to minimize visual differences between the two control strategies. **(B)** Biomimetic and arbitrary users calibrated their EMG controller so that specific biological hand gestures would engage specific bionic hand gestures (for the biomimetic strategy, these were matched). **(C)** Experimental design for the trained groups. The left column depicts examples of the training tasks included in a daily training session (see Supp. Video 1). The middle column depicts the timeline for each of the study visits. The right column depicts when the bionic hand gestures were introduced to participants in their training (on day 1 (D1): open and close; D2: pinch; D3: tripod). In the post-training generalization session, all participants (including the untrained participant group) learned to control the hand using a new set of hand gestures (i.e., new mapping). D=day.

## Results

To compare biomimetic and arbitrary control strategies, we tested 5 key features of bionic hand skill learning: (i) sense of embodiment, (ii) early training performance, (iii) late training performance, (iv) control automaticity and (v) post-training generalization to a new control mapping. To quantify motor performance, we focused on three myoelectric control skills: speed, dexterity and gesture switching.

To ensure any differences in skill learning were not driven by intrinsic differences in motor ability, prior to training, we tested participants on a ballistic reaching task using either the bionic hand (not yet turned on) or their biological left hand. We observed that all three groups had similar pre-training motor ability when wearing the bionic hand (*F*_(2,54)_=0.009, *p*=0.991; BF_incl_=0.14), as well as with their biological hand (*F*_(2,54)_=1.012, *p*=0.370; BF_incl_=0.291; Supp. Figure 1). Next, to ensure that any differences in skill learning were not driven by differences between EMG classifier performance for the biomimetic and arbitrary control strategies, we tested classifier performance immediately following device calibration. We found that the classifier had high performance for all participants with no differences in accuracy between control strategies (biomimetic average classification accuracy: 93% ± 5%; arbitrary: 92% ± 7%; *W*=167.0, *p*=0.631; BF_10_=0.37; Supp. Figure 2).). Therefore, any group differences potentially observed in skill learning could not be attributed to intrinsic differences in motor ability or classifier performance between control strategies.

### Biomimetic and arbitrary control show similar increases in bionic hand embodiment

To compare the two control strategies, we first assessed changes in the perceived (phenomenological) sense of embodiment of the bionic hand. Before and after training, participants were asked to respond to statements related to key embodiment categories: body ownership (*“it seems like the robotic hand is part of my body”*), agency (“*it seems like I am in control of the robotic hand”*) and visual appearance (“*it seems like I am looking directly at my own hand, rather than a robotic hand”;* Figure 2A; for a list of all questionnaire statements see Supp. Table 1). Comparing pre-to post-training scores, all trained participants reported a significant increase of embodiment in body ownership (*W*=84.0, *p*<0.001), and agency (*W*=12.0, *p*<0.001), but not visual appearance (*W*=137.0, *p*=0.132; BF_10_=0.48; Figure 2B). As subjective reports are particularly malleable to task demands (*19*), we also compared the training group to the untrained group (responding to the statements one week apart). This allowed us to confirm increased embodiment (post-minus pre-training ratings) in the trained groups relative to the untrained group [body ownership: *W*=263.0, *p*=0.045; agency: *W*=81.0, *p*<0.001; visual appearance: *W*=169.50, p<0.001; Figure 2C]. Importantly, we did not find differences between biomimetic and arbitrary users in the magnitude of this increase in embodiment reports, with the biomimetic/arbitrary group showing qualitatively (though not significantly) greater increase for ownership/agency, respectively [body ownership: *W*=266.50, *p*=0.143; BF_10_=0.53; agency: *W*=193.50, *p*=0.675; BF_10_=0.35; note that values are not corrected for multiple comparisons]. Overall, contrary to the common assumptions of biomimetic design, biomimetic control didn’t provide an increased sense of embodiment. Given no differences in perceived embodiment, this raised the interesting question whether we might observe differences between groups in skill learning.

**Figure 2.**
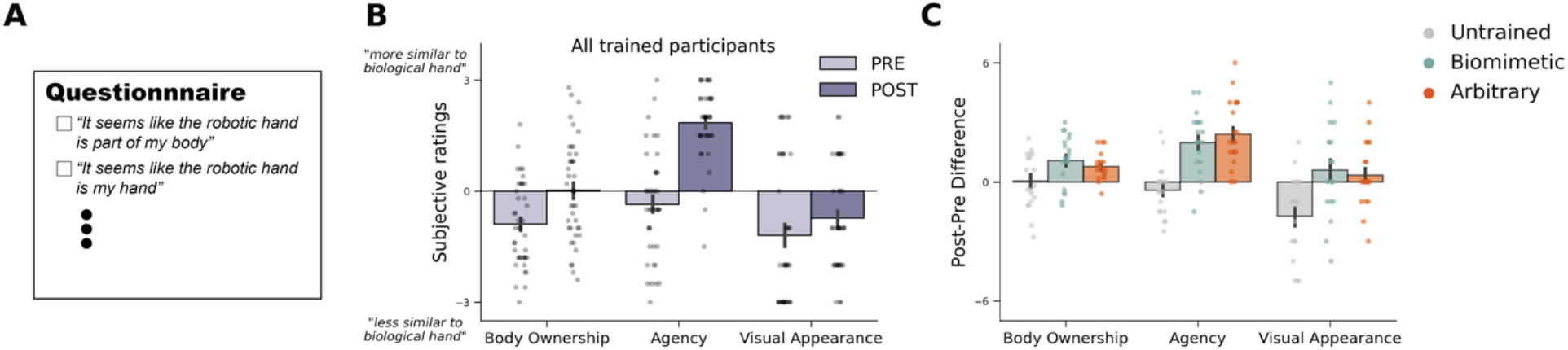
Trained participants show increased sense of bionic hand embodiment, regardless of control strategy. **(A)** An example item from the questionnaire used, designed to probe subjective sense of embodiment (all statements listed in Supp. Table 1). Scores were averaged over items of the same category (e.g., all body ownership questions averaged into a single value). **(B)** After training, trained participants showed significant increases in their sense of bionic hand embodiment on statements reflecting body ownership and agency. **(C)** Trained participants showed significant increases in embodiment scores compared to untrained participants on all categories, regardless of the control strategy users trained with. There were no significant differences between biomimetic and arbitrary users. Circles depict individual subject means (across relevant items). Values indicate group means ± standard error.

### Biomimetic control provides some early training speed benefits

We next focused on early training performance. To measure control speed, we quantified the ability to operate the hand using completion time on a modified version of the Southampton Hand Assessment Procedure (SHAP; Supp. Video 1). During the first training day, all participants were able to successfully complete the task, but biomimetic users were faster than arbitrary users (Figure 3A; D1 performance: *W*=85.0, *p*=0.001). Next, to quantify dexterity, we used the virtual eggs test which measures a users’ability to gently grasp and transport fragile (magnet-fused) blocks (i.e., ‘eggs’) without dropping or breaking them (Supp. Video 1). During the first training day, the majority of participants (75%) could not successfully transfer one egg without breaking it within the allocated time and there were no group differences (Figure 3B; D1 performance: *W*=185.50, *p*=0.897; BF_10_=0.34). To quantify the ability to switch between bionic hand gestures (close and pinch), we used completion time on a block stacking task that required participants to successfully grasp and transfer objects using the bionic hand close and pinch gestures, switching back and forth (Supp. Video 1). This ability could only be first tested on D2 because participants were only then trained on the second grasping gesture (pinch), thus providing gesture switching functionality. During the first attempt of this task, we observed no group differences in performance (Figure 3C; D2 performance: *W*=223.0, *p*=0.537; BF_10_=0.37). Overall, these findings suggest that biomimetic control affords early training benefits in speed, but not for dexterity and gesture switching.

**Figure 3.**
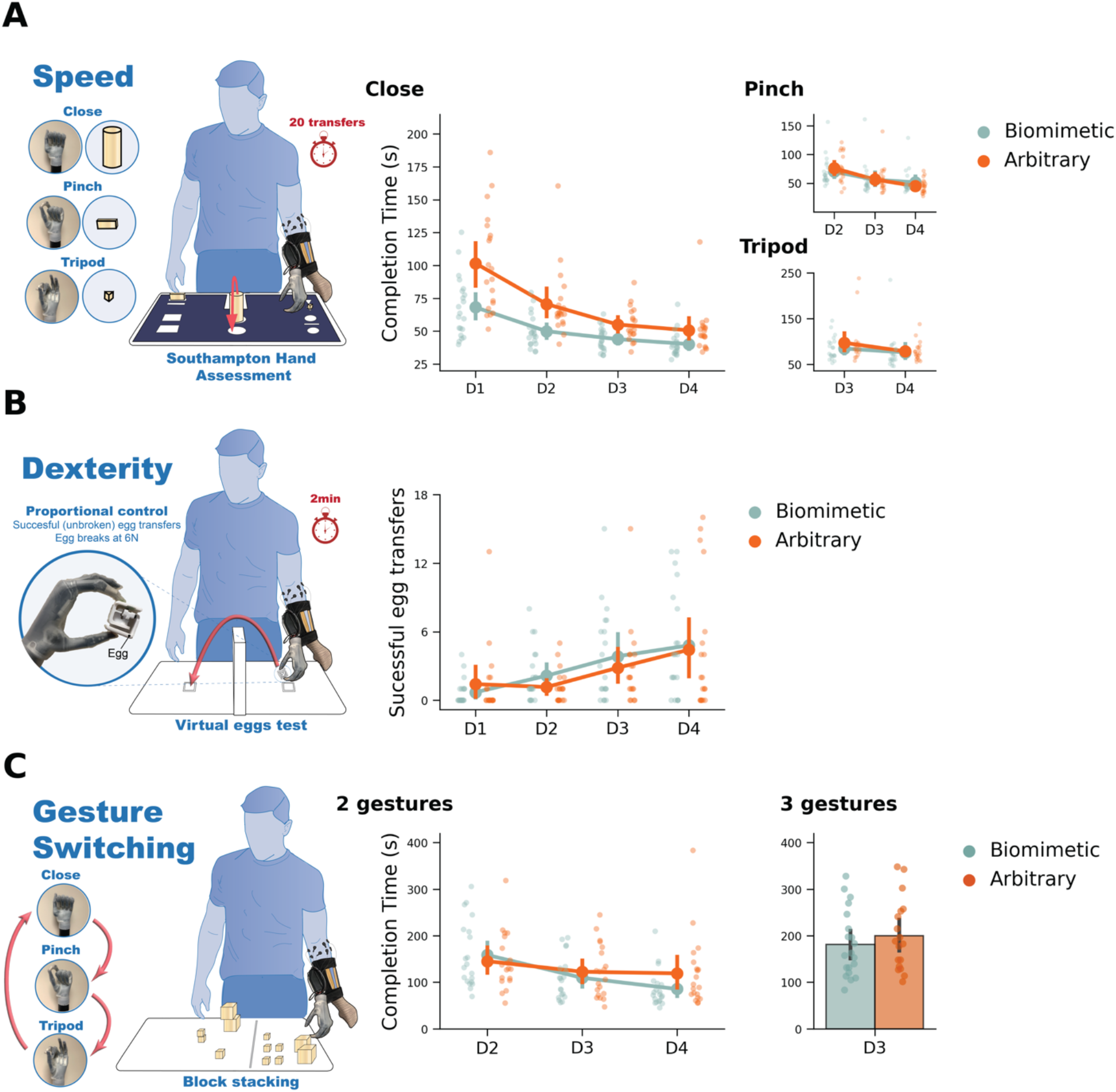
Bionic hand skill learning on speed, dexterity and gesture switching tasks. **(A)** Trained participants improved in control speed on all gestures. For the close gesture, biomimetic control was faster than arbitrary control across training sessions. **(B-C)** Trained participants improved in control dexterity (number of successful/unbroken egg transfers) and gesture switching across training sessions, regardless of control strategy. For gesture switching, because participants were trained on new grasping gestures each training day, we used two versions of this task. The 2 gestures version required successfully switching between close and pinch bionic gestures. The 3 gestures version required successfully switching between close, pinch and tripod bionic gestures. No significant differences were found between control strategies. See Supp. Video 1 for examples of all tasks. All other annotations are the same as described in Figure 2.

### Biomimetic advantage reduces with more training

Next, we examined late training motor performance (i.e., across the subsequent training days). For speed, on the SHAP, biomimetic users continued to outperform arbitrary users. All trained participants continued to improve performance with training (Figure 3A; main effect of *day: F*_(3,114)_=39.526, *p*<0.001). While the group differences narrowed with training (significant interaction between *day*group: F*_(3,114)_=5.659, *p*=0.001), biomimetic users were still faster than arbitrary users even in the last day of training (D4) (*W*=96.0, *p*=0.004). We also tested users on a different version of the SHAP that tests speed when using the pinch and tripod gestures. For these tasks, we observed significant improvements for all trained participants (Figure 3A; main effect of *day*; pinch: *F*_(2,78)_=48.435, *p*<0.001; tripod: *F*_(1,39)_=6.517, *p*=0.015), but no group differences on either task [main effect of *group;* pinch: *F*_(1,39)_=1.5e^-5^, *p*=0.997; BF_incl_=0.67; tripod: *F*_(1,39)_=0.676, *p*=0.416; BF_incl_=0.38; for all reported comparisons see Supp. Table 2].

For dexterity, we did not observe any group differences emerging with training. All trained participants improved in successful egg transfers across the training days (Figure 3B; main effect of *day*: *F*_(3,111)_=14.628, *p*<0.001) and there were no differences between groups (main effect of *group: F*_(1,37)_=0.233, *p*=0.632; BF_incl_=0.27; for all statistical comparisons see Supp. Table 2). On the last day of training (D4), both groups performed similarly (*W*=209.0, *p*=0.598; BF_10_=0.32).

Similarly, for gesture switching (i.e., switching between 2 grasping gestures: close, pinch), all trained participants improved in gesture switching speed [Figure 3C; main effect of *day*: *F*_(2,76)_=13.766, *p*<0.001] and there were no significant differences on average performance between groups (main effect of *group: F*_(1,38)_=0.044, *p*=0.835; BF_incl_=0.63). Additionally, when all gestures (3 grasping gestures: close, pinch, tripod) were incorporated into the task, we found no differences between groups (Figure 3D; *W*=172.0, *p*=0.646; BF_10_=0.34).

Overall, we observed a speed advantage for biomimetic users. However, this advantage was only observed for the close gesture (and its specific version of the SHAP) and the advantage was seen to reduce with training. Additionally, biomimetic control did not show any advantages, relative to the arbitrary, when learning dexterity and gesture switching. Instead, we found that training led to improvements, regardless of the control strategy.

### Biomimetic control provides more automatic control early in training

Another key component for successful integration with a bionic limb is the ability to multitask, such that attentional resources (e.g. focused exclusively on online control and movement planning) can be diverted towards other tasks without interfering with device control (i.e., control automaticity). On the first and last days of training, we tested the impact increased cognitive load would have on performance. The task required participants to perform arithmetic operations while simultaneously using the bionic hand to stack blocks (Supp. Video 1). To quantify the impact of increased cognitive load, we compared the number of blocks stacked with the counting task versus without (baseline). For counting performance, we observed that both groups performed the task similarly (main effect of *group: F*_(1,38)_=0.025, *p*=0.874; BF_incl_=0.32). For motor performance, we observed significant differences between groups across days (interaction between *day*group: F*_(1,38)_=9.896, *p*=0.003; Figure 4A). Looking at the first and last days separately, we observed that, on the first day of training, arbitrary users were more cognitively impaired than biomimetic users (*W*=286.50, *p*=0.019). However, on the last day of training, both groups were similarly affected by the cognitive load (*W*=176.0, *p*=0.533; BF_10_=0.36), suggesting that biomimetic control is more automatic early in training compared to arbitrary control, but arbitrary control becomes just as intuitive with continued training.

**Figure 4.**
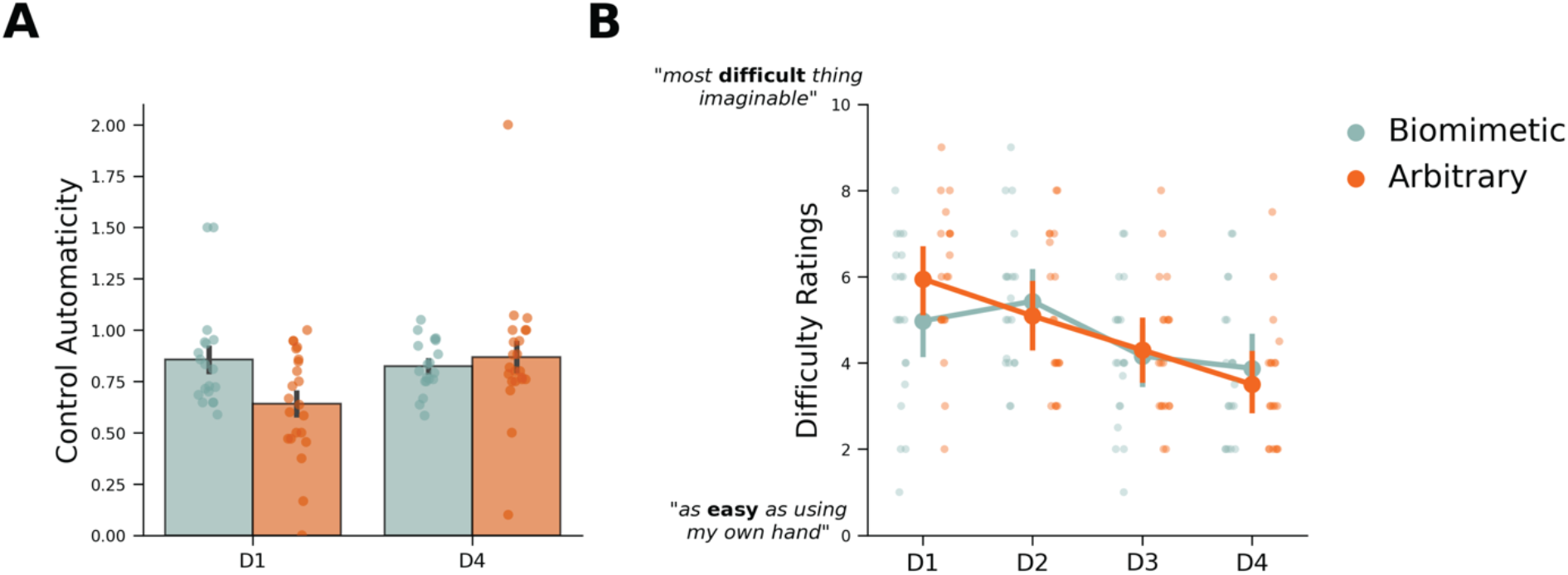
Arbitrary control is less automatic early in training but similar to biomimetic later. **(A)** Control automaticity was computed by dividing motor performance (number of blocks stacked) when simultaneously performing a counting task by motor performance alone. By this measure, higher values indicate less cognitive encumbrance, or more automatic control. Biomimetic control showed superior performance earlier in training, compared to arbitrary control. However, by the end of training, arbitrary control produced similar performance to that observed in the biomimetic group. **(B)** All trained participants reported control got easier across the training sessions, regardless of the control strategy. All other annotations are the same as described in Figure 2.

Another means of measuring automaticity is simply asking participants to rate their subjective sense of control difficulty at the end of every training day (see Methods). In these reports, all trained participants reported a significant decrease in control difficulty across days (Figure 4B; main effect of *day*: *F*_(3,114)_=21.298, *p*<0.001), but there were no average group differences in ratings across days (main effect of *group: F*_(1,38)_=0.041, *p*=0.840; BF_incl_=0.56).

Overall, we observed that biomimetic control provided increased control automaticity early in training compared to arbitrary. However, this advantage diminished when assessed later in training. Additionally, we did not find a significant impact of the control strategy on the subjective experience of control difficulty.

### Arbitrary strategy increases generalization to new control mappings

The ability to perform under a different set of conditions is a crucial aspect of prosthesis control. To test how well the learning for each control strategy generalizes to a new control mapping, we included a final post-training testing session where we re-calibrated the controller for all participants with a new set of gestures (see Methods and Figure 1C). All participants used the same set of gestures for this session. With this new control mapping, we tested users’ speed, dexterity and control difficulty. We, also, trained the untrained users to operate the hand with these gestures, providing a baseline for first-time use.

First, when testing speed on the SHAP in the generalization session, we found the biomimetic group matched performance of the untrained users (Figure 5A; *W*=202.0, *p*=0.534; BF_10_=0.38). Alternatively, the arbitrary group was significantly faster than untrained users (*W*=239.0, *p*=0.039). For trained participants, we also compared generalization speed performance to their first (D1) and last (D4) training day performance, when using their original control mappings. We observed a trend for significant group differences across days [D4 vs. generalization; interaction between *day*group: F*_(1,38)_=3.80, *p*=0.059]. Specifically, we found that biomimetic users speed dropped significantly between the two days (D4 vs. generalization: *W*=10.0, *p*<0.001), arbitrary users’ speed did not change (D4 vs. generalization: *W*=106.0, *p*=0.985; BF_10_=0.24).

**Figure 5.**
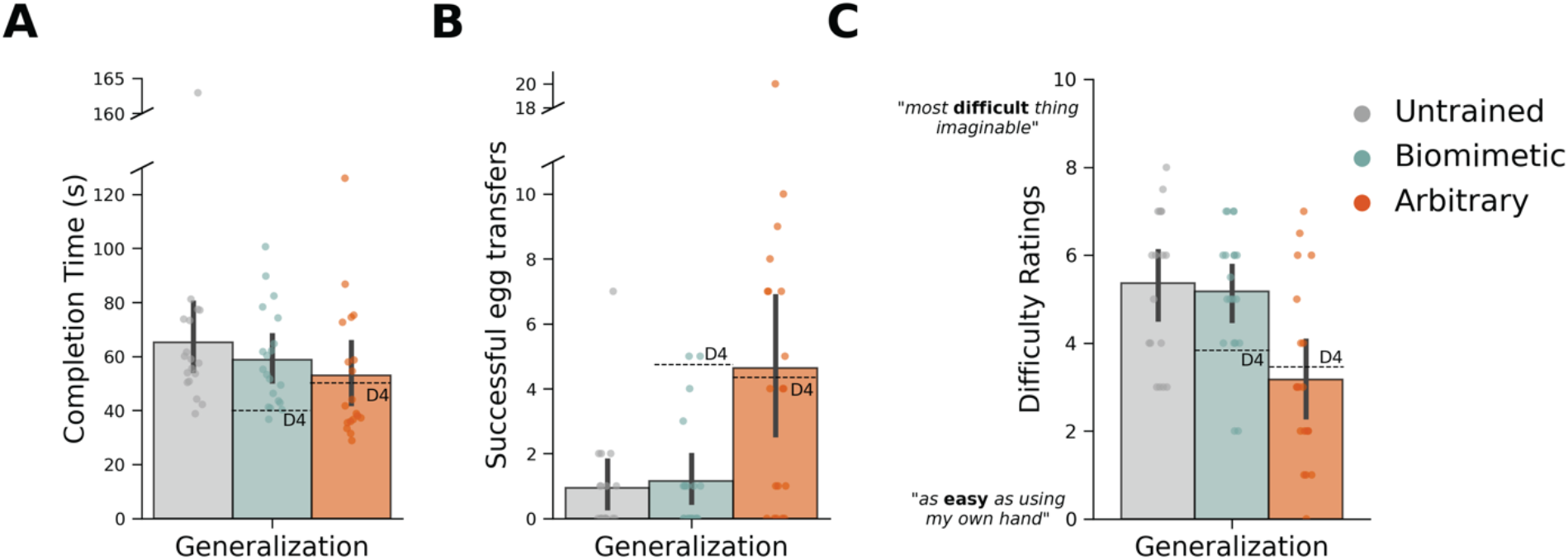
Arbitrary users show increased generalization. Performance on **(A)** speed, **(B)** dexterity and **(C)** control difficulty assessments for the generalization session. As a reference for previous performance, the dashed line denotes group mean performance on the last (D4) training session with their original control mappings. Biomimetic users showed significant impairments in comparison to performance during D4. Further, performance in the generalization tasks was similar between the biomimetic group and untrained participants using the bionic hand for the first time. Arbitrary users showed increased generalization such that performance with the new control strategy in the generalization session was similar to performance with the trained strategy on D4. All other annotations are the same as described in Figure 2.

Second, when testing dexterity in the generalization session, we, similarly, observed that biomimetic user performance matched the untrained group (Figure 5B; *W*=170.0, *p*=0.761; BF_10_=0.33), even returning to their pre-training (D1) performance level (*W*=17.50, *p*=0.328; BF_10_=0.35). Alternatively, the arbitrary group performed significantly better than both the biomimetic (*W*=104.50, *p*=0.013) and untrained groups (*W*=88.50, *p*=0.010). When comparing performance across days directly (D4 vs. generalization), we observed significant group differences in dexterity across days [interaction between *day*group: F*_(1,37)_=6.603, *p*=0.014]. Specifically, we found that biomimetic users’ dexterity dropped significantly between the two days (*W*=116.0, *p*=0.002). Alternatively, arbitrary users’ dexterity was the same as their previous training day (*W*=39.50, *p*=0.70; BF_10_=0.24), qualitatively even performing slightly better (D4 average eggs: 4.4; generalization average eggs: 4.6). In other words, while the biomimetic group showed no indication of generalization from the original to the new control mapping, the arbitrary strategy led to a full generalization of learning.

Finally, we observed that user’s subjective sense of control difficulty in the generalization session also supported these findings. We found that biomimetic users rated control difficulty in the generalization session similarly to the untrained group learning to operate the device for the very first time (Figure 5C; *W*=194.0, *p*=0.688; BF_10_=0.33), and even as difficult as biomimetic users’ first training day (D1 vs. generalization: *W*=72.50, *p*=0.867; BF_10_=0.25). Alternatively, arbitrary users reported control to be significantly easier than both the biomimetic (*W*=310.0, *p*=0.003) and untrained groups (*W*=287.50, *p*=0.002). Further, arbitrary users reported the difficulty to be as easy as their previous training day with their original control mapping (*W*=104.0, *p*=0.427; BF_10_=0.29) and qualitatively even slightly easier (D4 mean: 3.58 out of 10; generalization mean: 3.07 out of 10). When comparing responses across days directly (D4 vs. generalization), we observed significant differences in ratings between biomimetic and arbitrary groups [significant interaction between *day*group: F*_(1,38)_=6.857, *p*=0.013]. Specifically, we found that biomimetic user ratings were significantly more difficult in the generalization session compared to D4 (*W*=15.50, *p*=0.012). Alternatively, arbitrary user ratings were equally as difficult as their previous training day (*W*=104.0, *p*=0.427; BF_10_=0.27), qualitatively even slightly easier (D4 rating: 3.5; generalization rating: 3.1). Collectively, this demonstrates that arbitrary training provides increased control generalization to a novel control mapping.

## Discussion

It is a widely held assumption that control strategies designed to mimic the biological body might provide unique benefits to the user in terms of device learning, generalization, sense of embodiment, and automaticity (*5*, *8*, *11*, *20*–*27*). Contrary to this view, across a multitude of tasks, we observed few advantages for biomimetic control. We confirmed our predictions that biomimetic control was more intuitive for users, particularly in the early stages of learning (based on the cognitive load and speed tasks). However, when task difficulty was increased (more complex gestures, dexterous grasping and gesture switching), these advantages were mostly absent when compared to arbitrary users. Further, we observed that arbitrary users showed increased generalization to a new control mapping, while biomimetic users showed less capacity to generalize, performing similar to untrained participants. In addition, users subjective experience of perceived embodiment and control difficulty was not impacted by control strategy. Collectively, our findings provide a more balanced perspective of the neurocognitive challenges and opportunities of biomimetic and non-biomimetic control strategies. By challenging some of the core assumptions underlying biomimetic inspired design, our findings open up the potential for non-biomimetic control solutions for users.

Our findings are consistent with anecdotal evidence suggesting that arbitrary control approaches can be viable and even advantageous for artificial limb control. For example, the previous prosthetic hand winners at the Cybathalon – the Olympics for bionic technology – used devices that were explicitly designed to maximize functionality over biomimicry [2016 winner: Bob Radoc (pilot), DIPO Power Team (designers), Grip 5 Prehensor hand (device) and 2020 winner: Krunoslav Mihic (pilot), Andrj Đukić (designer), Maker Hand (device) (*28*, *29*)]. Additionally, on virtual tasks, research in amputees has shown that arbitrary myoelectric control in amputees can be learned, with one study showing amputees can learn to control an 8-target arbitrary myoelectric interface (*30*, *31*). Other studies have highlighted the emergence of arbitrary muscle synergies for EMG control (*32*, *33*). Most recently, new evidence supporting the versatility of non-biomimetic control has come from the field of motor augmentation. For example, people are able to rapidly learn to use a new body part (an extra robotic thumb) using 2 degrees of freedom operated via the users’ toes (*34*, *35*). Finally, efforts to prioritize functionality over biomimicry has led to the development of multiple creative and compelling control schemes for prosthetics (e.g., the use of footswitches, body powered devices, harnessing linear potentiometers, inertial measurement units to measure arbitrary gestures, RFID tags, proximity switches, co-contraction switches, etc.). Collectively, this evidence highlights the immense promise of non-biomimetic control solutions for assistive bionics.

Why did arbitrary control provide comparable degree of skill, and even outperform biomimetic control on generalization? Paradoxically, this could be a result of the cognitively challenging task demands related to adopting atypical gestures for grasp control. Our ability to skillfully use our hands is developed over a lifetime of experience, in which information on motor synergies, object knowledge and action semantics is meticulously weaved to construct efficient motor command pipelines (*36*). Arbitrary users have to learn to associate hand motor commands with entirely unrelated action goals (previously associated with different motor commands). This introduces high amounts of contextual novelty into arbitrary user’s skill learning. While early in training, this might be disadvantageous (as demonstrated by arbitrary control’s increased cognitive demands relative to the biomimetic group), there is ample evidence to suggest that this could actually improve overall learning. For example, learning with a more varied input that increases task difficulty is initially slower, but typically yields better generalization [for a review see (*37*)]. Accordingly, research has demonstrated that more complex associations end up being learned more robustly and are better retained, than easy-to-learn associations (*38*). One potential explanation for this is that easy-to-learn associations (in our study, biomimetic control) rely more heavily on working memory which allows for very fast learning of information that is not durably retained, while learning complex associations relies on reinforcement learning which allows for slow, integrative learning of associations that is robustly stored. In summary, the contextual differences between the strategies and the recruitment of reinforcement learning for arbitrary learning may potentially explain why arbitrary control outperformed biomimetic when generalizing to a novel control mapping.

The considerations above suggest why arbitrary strategies can be beneficial, but we must also consider the alternative perspective – that is – could biomimetic control be disadvantageous? There are several potential explanations why biomimicry could have disadvantages, primarily due to its’ ambition to stay so close to the body. While modern bionic limbs are increasingly biomimetic in design and control, they are not at all the same as biological limbs. Modern prosthetics have speed delays, limited dexterity, and functionality, making their operation a lot clumsier than a biological hand. Devices are generally heavy, not particularly durable and their sensory feedback is impoverished. These discrepancies between biological and bionic limb control may be promoting an ‘Uncanny Valley’-type phenomenon for users (*39*).The Uncanny Valley proposes that an individual’s response to a humanoid robot shifts from empathy to repulsion as the humanoid approaches, but fails to achieve, humanlike appearance [e.g., Tom Hanks in the Polar Express (*40*); the humanoid horror doll in M3gan (*41*)]. While this phenomenon is often used in relation to how we *see* artificial bodies, the framework also seems relevant to how we *control* artificial bodies. As device control become more and more biomimetic, but not capable of reaching true biomimicry, it may be creating a more active neurocognitive competition between the priors, sensory predictions, and motor commands for how amputees controlled their pre-existing limb and how they represent and control their prosthetic limb. Therefore, non-biomimetic control solutions may be more beneficial because the motor control plan can be developed from scratch (i.e., independent of pre-existing biological limb control) and therefore can avoid any potential conflict.

It is important to note that our experimental design may have potential limitations. First, due to the limited state of modern prosthetic technology, the biomimetic strategy we implemented was not truly pure biomimetic (i.e., the same as biological hand control). However, if we consider biomimicry as a spectrum of strategies closer and further away from the biological body, the biomimetic control we tested falls closer to biological control than the arbitrary control. Second, our training was restricted to only 4 daily lab-based sessions, and it is possible that with additional training more differences would have emerged between the groups. While this might seem like limited training, a recent survey reported that about half (43%) of amputees that were trained to use a prosthesis received between just 1 to 3 training visits with their device (*16*). Furthermore, the arbitrary gestures we have tested involved simple finger extensions, and it is likely that more careful curation of control mapping could have produced much greater benefits for the arbitrary control. For these potential limitations, in our view, the fact that arbitrary users matched biomimetic performance after such minimal training is promising. A further limitation is that we tested able-bodied participants instead of limbless participants, the clinical population this research will impact. However, considering the primary aim of the study was to mimic bionic limb control to the biological body, we thought it was necessary for us to have direct access to participant’s biological limb to ensure true biomimicry. Moreover, research is now mounting to indicate that amputation might not induce far reaching changes to the motor representation of the missing limb (*42*–*44*), even with regards to motor learning (*45*). Regardless, future studies should continue to evaluate personalized control solutions for acquired and congenital limbless individuals.

In summary, due to the current limitations in modern prosthetic technologies not yet matching those available to Luke Skywalker, researchers and engineers should continue exploring both biomimetic and non-biomimetic control solutions. While biomimetic design is an understandable starting point when designing human-machine interfaces, it should not necessarily be the ultimate, end-all goal. From a cognitive neuroscience perspective, there are multiple considerations for and against biomimetic control. So, practically speaking, how biomimetic should we go? Based on our findings, this depends entirely on the purpose of the prosthesis. If the device is intended for short-term use, simple functionality, or where training opportunities are limited, then biomimetic-inspired control options make a lot of sense. However, if the purpose is to design versatile devices with multiple functions for long-term use, at least with modern EMG pattern recognition technology, non-biomimetic control solutions may provide a useful means to enhance certain aspects of bionic hand motor learning. Further, abandoning the unrealistic ambition of true biomimicry opens up endless possibilities for users and engineers to develop a variety of different control solutions. We suggest that engineers and prosthetists involved in the commercial and clinical delivery of this technology should prioritize flexibility – educating users on the spectrum of biomimetic-to-arbitrary control strategies available to them such that personalized, user-specific control strategies can be selected based on individual user requirements. In our experience, when users are educated with the knowledge and confidence that multiple control approaches are possible, there is a higher likelihood that devices will meet user expectations and requirements. Personalized control strategies will help to propel the industry closer to the actual goal: more satisfied prosthesis users.

## Supporting information

Supplementary Video 1. Video examples of the tasks.

## Supplementary Figures and Tables

**Supplementary Figure 1.**
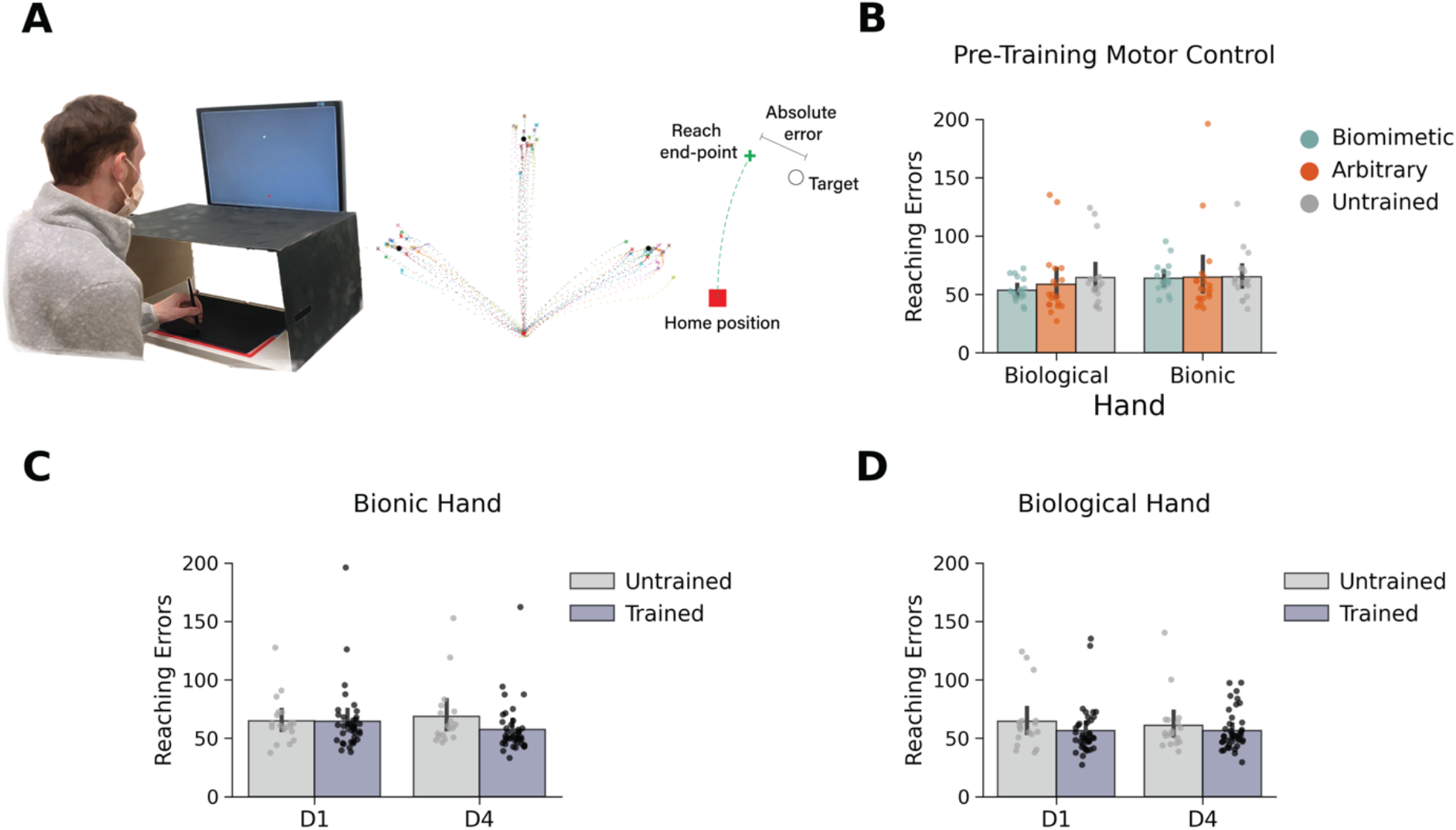
Similar motor abilities between groups when using biological and bionic hands. **(A)** A participant performing ballistic reaches (with no corrective movements) to virtual targets using a digitizing tablet and stylus. The task was performed with either participant’s left biological hand or the bionic hand locked around the stylus (see Methods for a description). Not shown in the figure image, during the task, all participants wore a barber cape over the apparatus to remove any potential visual feedback of their arm/hand. Participants performed 60 reaches with each hand (biological and bionic) to 3 different virtual targets (example of a participant reaches shown in panel A). The primary measure of motor ability we quantified is the average absolute error between reach end points and the virtual target location. **(B)** Before training, all groups made similar reaching errors when reaching with their biological hand or the bionic hand. **(C)** After-training, trained participants (biomimetic and arbitrary combined) made smaller reaching errors, on average, than untrained participants when using the bionic hand (*W*=498.0, *p*=0.020), but similar errors before training (*W*=397.0, *p*=0.551; BF_10_=0.29). **(D)** When using their biological hand, trained and untrained participants showed similar reaching errors both before (*W*=444.0, *p*=0.164; BF_10_=0.45) and after training (*W*=409.0, *p*=0.418; BF_10_=0.44). Circles depict individual subject means (across relevant items). Values indicate group means ±standard error.

**Supplementary Figure 2.**
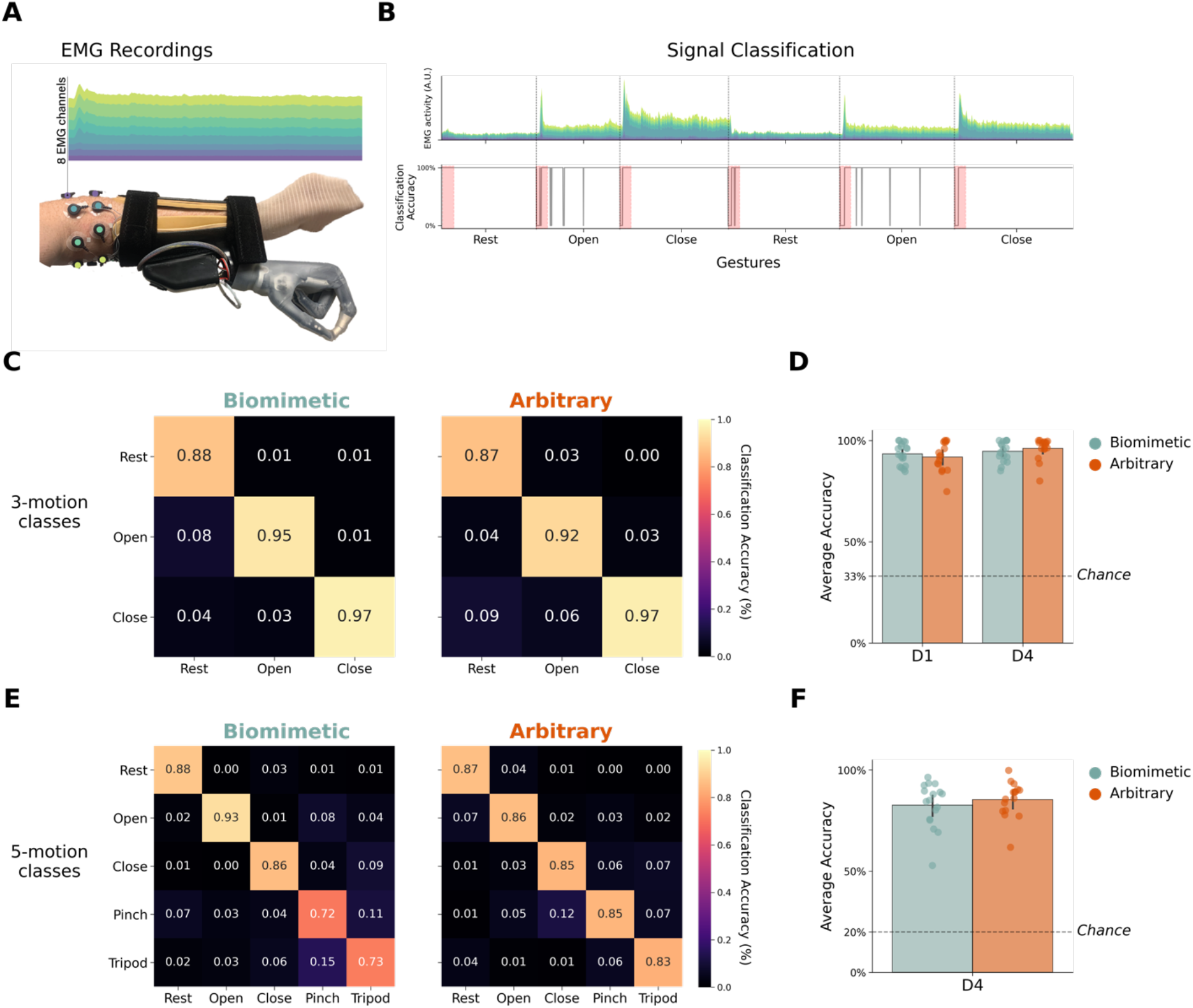
Similar EMG classification accuracy for biomimetic and arbitrary control strategies. **(A)** EMG-signals were acquired from the bionic hand system and classified by the Coapt EMG controller. **(B)** To measure classification accuracy, we asked participants to engage and hold each of the calibrated hand gestures for 20 seconds each (see Methods). During this assessment, the Coapt controller outputted a real-time classification decision (updated every 50ms) for which gesture was being performed. Comparing this output to the gesture participants were instructed to perform by the experimenter, we computed the classification accuracy for each gesture. The time-window of interest was approximately the first 2.5 seconds of each 20 second gesture trial (shown in panel B as a red window). The rationale for this time-window was to roughly capture the control experience during functional use. The data plotted depicts an example subject’s EMG activity and controller classification during the task. **(C).** Group average classification accuracy matrices for 3-gesture classes (rest, open close) groups assessed on D1, immediately following controller calibration. **(D)** Both trained groups showed a significant increase in average classification accuracy (main effect of day: *F*_(1,31)_=5.647, *p*=0.024) between the first (D1) and last day of training (D4). No significant differences were found between the two training strategies before training (*W*=167.0, *p*=0.631; BF_10_=0.37) or after training (*W*=121.50, *p*=0.437; BF_10_=0.37). **(E)** Group average classification accuracy matrices for 5-gesture classes (rest, open close, pinch, tripod), assessed on D4, at the end of training. (F) No differences between trained groups in average classification accuracy (average of the 5-motion-class diagonal; *W*=117.50, *p*=0.517; BF_10_=0.42). Circles depict individual subject means (across relevant items). Values indicate group means ±standard error.

**Supplementary Figure 3.**
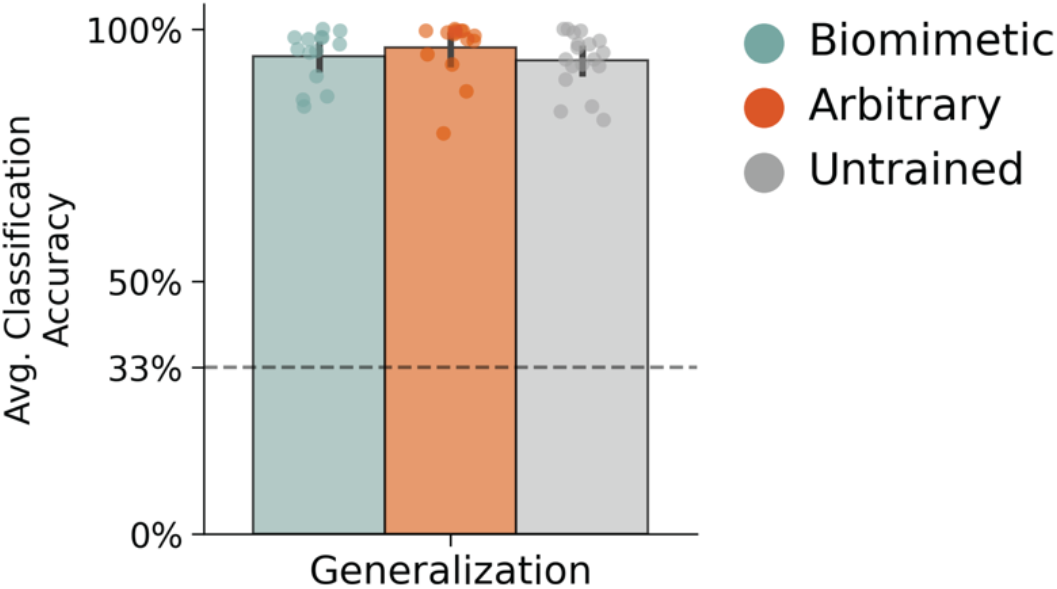
Similar classification accuracy for all groups during generalization session. Average classification accuracy for the generalization control mappings was calculated in the same way as Supp. Figure 2.

**Supplementary Video 1. Video examples of the tasks.**

**Supplementary Table 1.** Embodiment questionnaire statements divided into three categories.

|  |
| --- |
| **Body Ownership** |
| 1. "It seems like the robotic hand belongs to me" |
| 2. "It seems like the robotic hand is my hand" |
| 3. "It seems like the robotic hand is part of my body" |
| 4. "It feels like my robotic hand is a foreign body" |
| 5. "It feels like my robotic hand is fused with my body" |
| 6. "It seems like I have three hands" |
| **Agency** |
| 1. "It seems like I can move the robotic hand fingers if I want" |
| 2. "It seems like I am in control of the robotic hand" |
| **Body image** |
| 1. "It seems like I am looking directly at my own hand, rather than a robotic hand" |

**Supplementary Table 2.** All statistical analyses.

| Days | Condition - Measure | Statistical Test | Result |
| --- | --- | --- | --- |
| **Speed – Southampton Hand Assessment Procedure** |  |  |  |
| D1,D2,D3,D4 | Object1 – Completion Time | RmANOVA:<br>Days: 4<br>Groups: 2<br>Mann Whitney<br>Bayes Mann-Whitney | Day x Group:<br>$F_{(3,114)}=6.892$, $p<0.001$<br>Day:<br>$F_{(3,114)}=67.970$, $p<0.001$<br>Group:<br>$F_{(1,38)}=14.031$, $p<0.001$<br>D1: $W=85.0$, $p=0.001$<br>D2: $W=67.0$, $p<0.001$<br>D3: $W=93.50$, $p=0.004$<br>D4: $W=96.0$, $p=0.004$<br>D1: $BF_{10}=15.961$<br>D2: $BF_{10}=20.952$<br>D3: $BF_{10}=8.145$<br>D4: $BF_{10}=4.938$ |
| Generalization | Object1 – Completion Time | ANOVA:<br>Groups: 3<br>Mann Whitney<br>Bayes Mann-Whitney | $F_{(2,54)} = 1.212$, $p = .306$<br>Ctrl to Bio: $W=202.0$, $p=0.534$<br>Ctrl to Arb: $W=239.0$, $p=0.039$<br>Bio to Arb: $W=250.0$, $p=0.095$<br>Ctrl to Bio: $BF_{10} = .383$<br>Ctrl to Arb: $BF_{10} = 1.505$<br>Bio to Arb: $BF_{10} = .873$ |
| D2,D3,D4 | Object2 – Completion Time | RmANOVA:<br>Days: 3<br>Groups: 2<br>Mann Whitney<br>Bayes Mann-Whitney | Day x Group:<br>$F_{(2,76)} = 3.491$, $p=0.035$<br>Day:<br>$F_{(2,76)} = 47.583$, $p < .001$<br>Group:<br>$F_{(1,38)} = 1.087$, $p = .304$<br>D2: $W=263.0$, $p=0.088$<br>D3: $W=232.0$, $p=0.390$<br>D4: $W=175.0$, $p=0.524$<br>D2: $BF_{10}=0.722$<br>D3: $BF_{10}=0.384$<br>D4: $BF_{10}=0.453$ |
| D3,D4 | Object3 – Completion Time | RmANOVA:<br>Days: 2<br>Groups: 2<br>Mann Whitney<br>Bayes Mann-Whitney | Day x Group:<br>$F_{(1,38)}=0.375$, $p=0.544$<br>Day:<br>$F_{(1,38)}=5.196$, $p=0.028$<br>Group:<br>$F_{(1,38)}=0.185$, $p=0.670$<br>D3: $W=177.0$, $p=0.555$<br>D4: $W=158.0$, $p=0.270$<br>D3: $BF_{10}=0.362$<br>D4: $BF_{10}=0.472$ |
| **Dexterity - Virtual eggs test** |  |  |  |
| D1, D2,D3,D4 | Number of successful (unbroken) eggs transferred | RmANOVA:<br>Days: 4<br>Groups:2<br>Mann Whitney<br>Bayes Mann-Whitney | Day X Group:<br>$F_{(3,111)} = 0.921$, $p = .433$<br>Day:<br>$F_{(3,111)} = 14.628$, $p < .001$<br>Group:<br>$F_{(1,37)} = .233$, $p = .632$<br>D1: $W=185.50$, $p=0.897$<br>D2: $W=239.50$, $p=0.154$<br>D3: $W=216.0$, $p=0.468$<br>D4: $W=209.0$, $p=0.598$<br>D1: $BF_{10}=0.349$<br>D2: $BF_{10}=0.648$<br>D3: $BF_{10}=0.389$<br>D4: $BF_{10}=0.328$ |
| Generalization | Number of successful (unbroken) eggs transferred | ANOVA:<br>Groups: 3<br>Bayesian ANOVA<br>Mann Whitney<br>Bayes Mann-Whitney | $F_{(2,54)} = 7.787$, $p = 0.001$<br>$BF_{10}=32.417$<br>Ctrl to Arb: $W=88.50$, $p=0.010$<br>Ctrl to Bio: $W=170.0$, $p=0.716$<br>Bio to Arb: $W=104.50$, $p=0.013$<br>Ctrl to Bio: $BF_{10}=0.351$<br>Ctrl to Arb: $BF_{10}=2.530$<br>Bio to Arb: $BF_{10}=2.521$ |
| **Gesture Switching – Block stacking** |  |  |  |
| D2,D3,D4 | 2 gesture version – completion time | RmANOVA:<br>Days: 3<br>Groups: 2<br>Mann Whitney<br>Bayes Mann<br>Whitney | Day X Group:<br>$F_{(2,76)}=3.602, p=0.032$<br>Day:<br>$F_{(2,76)}=13.766, p<0.001$<br>Group:<br>$F_{(1,38)}=0.044, p=0.835$<br>D2: $W=223.0, p=0.537$<br>D3: $W=194.0, p=0.689$<br>D4: $W=154.0, p=0.149$<br>D2: $BF_{10}=0.379$<br>D3: $BF_{10}=0.319$<br>D4: $BF_{10}=0.844$ |
| D3 | 3 gesture version – completion time | Mann Whitney<br>Bayes Mann-Whitney | $W=172.0, p=0.646$<br>$BF_{10}=0.349$ |
| **Control Automaticity - Cognitive Load** |  |  |  |
| D1,D4 | Control Automaticity (blocks stacked with load divided by blocks stacked without load) | RmANOVA:<br>Days: 2<br>Groups: 2<br>Mann Whitney<br>Bayes Mann-Whitney | Day X Group:<br>$F_{(1,38)}=9.896, p=0.003$<br>Day:<br>$F_{(1,38)}=5.475, p=0.022$<br>Group:<br>$F_{(1,38)}=1.753, p=0.193$<br>D1: $W=286.50, p=0.019$<br>D4: $W=176.0, p=0.533$<br>D1: $BF_{10}=286.50$<br>D4: $BF_{10}=0.365$ |
| **Control Automaticity - Subjective Control Difficulty** |  |  |  |
| D1, D2, D3, D4 | Raw scores | RmANOVA:<br>Days: 4<br>Groups: 2<br>Mann Whitney<br>Bayes Mann-Whitney | Day X Group:<br>$F_{(3,114)}=2.924, p=0.037$<br>Day:<br>$F_{(3,114)}=21.298, p<0.001$<br>Group:<br>$F_{(1,38)}=0.041, p=0.840$<br>D1: $W=156.0, p=0.159$<br>D2: $W=242.50, p=0.397$<br>D3: $W=197.50, p=0.956$<br>D4: $W=232.0, p=0.566$<br>D1: $BF_{10}=0.824$<br>D2: $BF_{10}=0.382$<br>D3: $BF_{10}=0.312$<br>D4: $BF_{10}=0.377$ |
| D4, Generalization | Raw scores | RmANOVA:<br>Days: 2<br>Groups: 2<br>Mann Whitney<br>Bayes Mann-Whitney<br>Wilcoxon signed-<br>rank<br>Bayes Wilcoxon | Day X Group:<br>$F_{(1,38)}=6.857, p=0.013$<br>Day:<br>$F_{(1,38)}=2.469, p=0.124$<br>Group:<br>$F_{(1,38)}=7.626, p=0.009$<br>D4: $W=232.0, p=0.566$<br>Generalization: $W=310.0, p=0.003$<br>D4: $BF_{10}=0.377$<br>Generalization: $BF_{10}=30.395$<br>Bio: D4 vs. Gen: $W=15.50, p=0.012$<br>Arb: D4 vs. Gen: $W=104.0, p=0.427$<br>Bio: D4 vs. Gen: $BF_{10}=9.539$<br>Arb: D4 vs. Gen: $BF_{10}=0.279$ |
| Generalization | Raw scores | ANOVA:<br>Groups: 3<br>Mann Whitney<br>Bayes Mann-Whitney | $F_{(2,55)}=9.178, p<.001$<br>Ctrl to Arb: $W=287.50, p=0.002$<br>Ctrl to Bio: $W=194.0, p=0.688$<br>Bio to Arb: $W=310.0, p=0.003$<br>Ctrl to Bio: $BF_{10}=0.331$<br>Ctrl to Arb: $BF_{10}=28.721$<br>Bio to Arb: $BF_{10}=30.395$ |
| **Sense of Embodiment** |  |  |  |
| Pre, Post | Raw scores | RmANOVA:<br>Days: 2<br>Groups: 2 (trained<br>vs. untrained) | <u>Body Ownership</u><br>Day X Group:<br>$F_{(1,58)}=7.621, p=0.008$<br>Day:<br>$F_{(1,58)}=9.158, p=0.004$<br>Group:<br>$F_{(1,58)}=1.359, p=0.248$ |
| | | Mann-Whitney<br>Bayes Mann-Whitney<br>Run for each category separately | Pre: $W=412.50$, $p=0.720$; $BF_{10}=0.291$<br>Post: $W=262.50$, $p=0.044$; $BF_{10}=1.861$<br><u>Agency</u><br>Day X Group:<br>$F_{(1,58)}=39.323$, $p<0.001$<br>Day:<br>$F_{(1,58)}=18.082$, $p<0.001$<br>Group:<br>$F_{(1,58)}=8.530$, $p=0.005$<br>Pre: $W=461.50$, $p=0.252$; $BF_{10}=0.581$<br>Post: $W=76.50$, $p<0.001$; $BF_{10}=124.245$<br><u>Visual Appearance</u><br>Day X Group:<br>$F_{(1,58)}=15.998$, $p<0.001$<br>Day:<br>$F_{(1,58)}=5.359$, $p=0.024$<br>Group:<br>$F_{(1,58)}=0.217$, $p=0.643$<br>Pre: $W=501.50$, $p=0.070$; $BF_{10}=0.978$<br>Post: $W=184.50$, $p<0.001$; $BF_{10}=34.825$ |
| Pre and Post | Post-Pre difference score | Mann Whitney Groups: 2 (trained vs. untrained)<br>Mann-Whitney Groups: 2 (bio vs. arb)<br>Bayes Mann-Whitney Groups: 2 (bio vs. arb) | Body ownership: $W=263.0$, $p=0.045$<br>Agency: $W=81.0$, $p<0.001$<br>Visual appearance: $W=169.50$, $p<0.001$<br>Body ownership: $W=266.50$, $p=0.143$<br>Agency: $W=193.50$, $p=0.675$<br>Visual appearance: $W=228.50$, $p=0.632$<br>Body Ownership: $BF_{10}=0.532$<br>Agency: $BF_{10}=0.385$ |
| **Classification Accuracy** |  |  |  |
| D1 | 3-motion-class (avg. of rest, open, close) | Mann Whitney<br>Bayes Mann-Whitney | $W=167.0$, $p=0.631$<br>$BF_{10}=0.375$ |
| D1, D4 | 3-motion-class (avg. of rest, open, close) | RmANOVA:<br>Days: 2<br>Groups: 2 (bio vs. arb)<br>Mann-Whitney<br>Bayes Mann-Whitney | Day X Group:<br>$F_{(1,31)}=2.622$, $p=0.116$<br>Day:<br>$F_{(1,31)}=5.647$, $p=0.024$<br>Group:<br>$F_{(1,31)}=0.001$, $p=0.970$<br>Pre: $W=167.0$, $p=0.631$<br>Post: $W=121.50$, $p=0.437$<br>Pre: $BF_{10}=0.375$<br>Post: $BF_{10}=0.371$ |
| D4 | 5-motion-class (avg. of rest, open, close, pinch, tripod) | Mann-Whitney<br>Bayes Mann-Whitney | $W=117.50$, $p=0.517$<br>$BF_{10}=0.422$ |
| Generalization | 3-motion-class (avg. of rest, open, close) | ANOVA:<br>Groups: 3 | $F_{(2,44)}=0.874$, $p=0.424$ |
| **Motor control** |  |  |  |
| Pre | Bionic hand - Absolute error | ANOVA:<br>Groups: 3<br>Bayesian ANOVA:<br>Groups: 3 | $F_{(2,54)}=0.009$, $p=0.991$<br>$BF_{10}=0.14$ |
| Pre | Biological hand - Absolute error | ANOVA:<br>Groups: 3<br>Bayesian ANOVA:<br>Groups: 3 | $F_{(2,54)}=1.012$, $p=0.370$<br>$BF_{10}=0.291$ |
| Pre and Post | Bionic hand – Absolute error | RmANOVA:<br>Days: 2<br>Groups: 2 (trained vs. untrained)<br>Mann Whitney | Day X Group:<br>$F_{(1,50)}=4.010$, $p=0.051$<br>Day:<br>$F_{(1,50)}=2.522$, $p=0.118$<br>Group:<br>$F_{(1,50)}=1.725$, $p=0.195$<br>Pre: $W=397.0$, $p=0.551$<br>Post: $W=498.0$, $p=0.020$ |

## Methods

The study and its experimental procedures were approved by the NIH Institutional Review Board (NCT00001360, 93M-0170). The study reported here was conducted as part of a larger experiment (see https://osf.io/3m592/ for pre-registration of the full experimental protocol). Below we only report the methodology relevant for the results detailed in the manuscript.

### Participants

Sixty-one healthy volunteers [40 females; mean age = 24.8 ±0.66; all right handed) were recruited from the National Institute of Health community and the Washington DC metro area and were randomly assigned to one of the following study groups: biomimetic (n = 21; 14 females; mean age 23.9 ±0.57), arbitrary (n = 21; 12 females; mean age 25.9 ±1.28) or untrained (n = 19; 14 females; mean age 24.6 ±1.41). All participants were unaware of the other participant groups to minimize any potential biases on participant performance. All participants had no known motor disorders. Criteria for participant inclusion were determined prior to data collection according to the study population guidelines approved by the NIH Institutional Review Board as a part of the study protocol (93-M-0170, NCT00001360). All participants gave their written informed consent before participating in the study and were compensated monetarily for their time.

Two additional participants were recruited, but not included in the present study due to incomplete datasets.

### Experimental design

To quantify bionic hand skill learning, we implemented a longitudinal experimental design (Figure 1C), involving 6 experimental sessions conducted across 6 days (1 session per day, within a 1-week period), as summarized in Figure 1. All trained participants (biomimetic and arbitrary groups) underwent (i) a familiarization session (2 hours), introducing the equipment and completing some pre-training motor control assessments; (ii) four training sessions (2-3 hours) conducted over four consecutive days (one session per day) and (iii) a final generalization behavioral session (2 hours).

Untrained participants underwent a modified schedule: (i) the familiarization session (2 hours) and (ii) the generalization behavioral session (1-week later; 2 hours). The generalization session was the first-time untrained participants were able to have active control over the device.

### Biomimetic, arbitrary and generalization control mappings

Biomimetic and arbitrary control mappings differed in the biological hand gestures that were required to engage the bionic hand classifier (Figure 1B). The biomimetic control mappings included: open hand (biological) = open hand (bionic), close hand (biological) = close hand (bionic), pinch (biological) = pinch (bionic), and tripod (biological) = tripod (bionic). The arbitrary control mappings included: 1 finger (biological) = open hand (bionic), 2 fingers (biological) = close hand (bionic), 3 fingers (biological) = pinch (bionic), and 4 fingers (biological) = tripod (bionic). Finally, during the generalization session, all participants (untrained participants included) learned a novel control strategy. The generalization control mappings included: thumbs-up (biological) = open hand (bionic) and thumb-pinky pinch (biological) = close hand (bionic). The generalization gestures were intended to be gestures that were a hybrid of the gestures in the biomimetic and arbitrary gestures. Additionally, our reasoning for having all groups use the same gestures in this session was that it would allow us to directly match performance across groups.

### Bionic hand setup and calibration

#### Bionic hand system and setup

A custom-made left-hand bionic system was created for this study (Figure 6). A custom laminated fiber glass socket was fitted around participant’s forearm. The socket includes a lamination ring and a coaxial plug (Ossur) which interfaces with an i-LIMB Quantum Hand QWD [OSSUR; (*46*)]. An i-limb skin Active glove (Ossur) was worn on the hand. A custom thermoplastic component housed a Coapt COMPLETE CONTROL Gen2 pattern recognition system [Coapt, LLC; firmware v1.27; software v.1.1.9 (*18*)] and rechargeable Lithium Polymer batteries (Ossur, model: 704374; battery rating: 7.4V, 2000mAh; capacity: 14.8Wh). The COMPLETE CONTROL system is a clinically available EMG pattern recognition system. The thermoplastic component was attached to the carbon fiber socket using Velcro. The socket was tightly positioned around participant’s forearm using Velcro straps. A custom EMG cable attached to the Coapt EMG controller and connected to electrodes on the upper-forearm.

**Figure 6.**
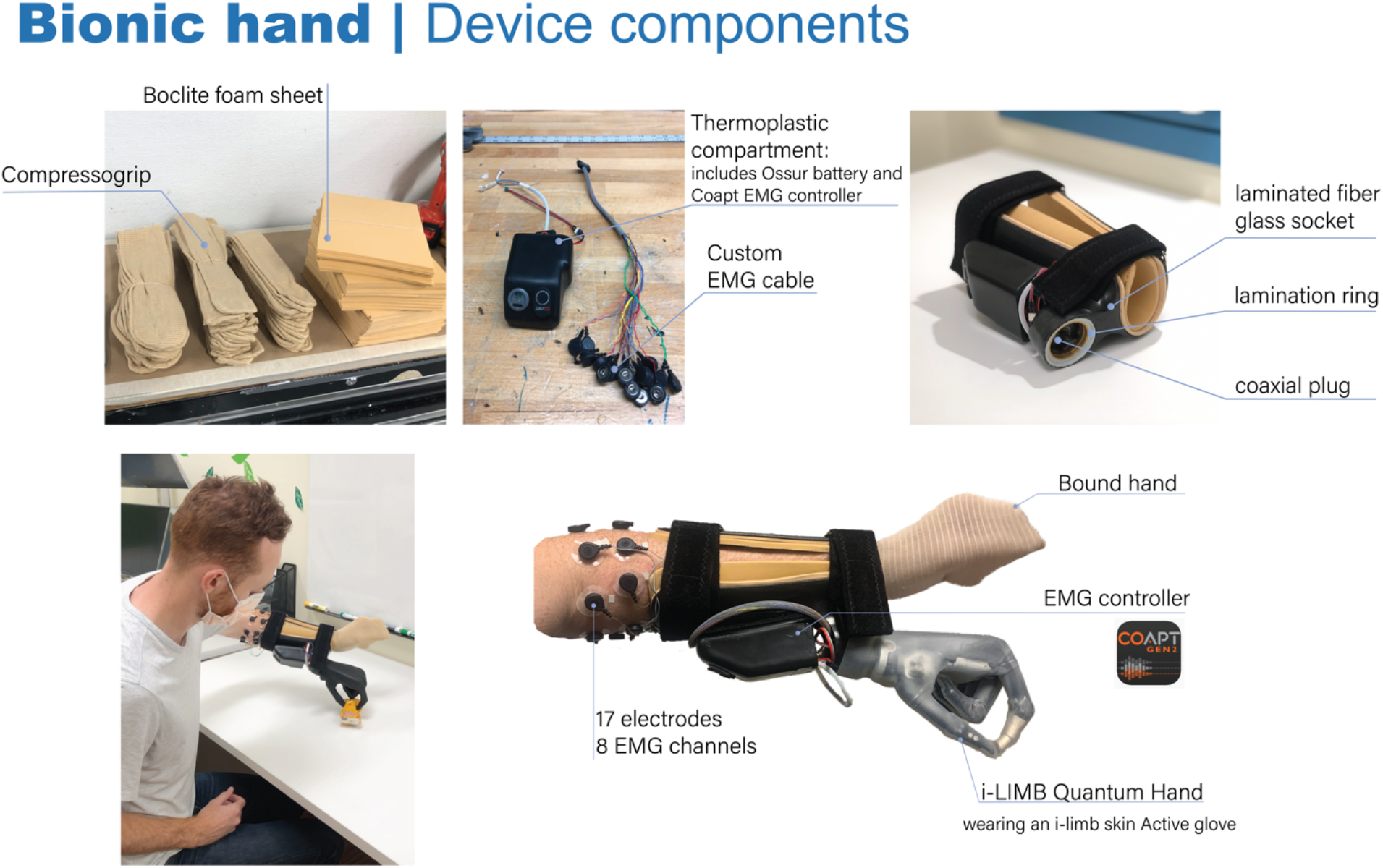
Bionic hand device components. An image breakdown of all of the individual components incorporated in the bionic hand system.

An Ossur-certified researcher fitted the bionic hand to each participant’s left biological arm. To reduce visual differences between groups, the biological hand was bound in an elastic, sewn fabric (Compressogrip). Participants then were fitted with a custom Boclite foam sheet (to reduce skin irritation). To ensure no interference with EMG signal, prior to study participation, all participants shaved the hair on the left forearm. Prior to electrode placement, the skin of the forearm was cleaned with water and a mildly abrasive paste. Once cleaned, eight EMG electrode pairs (bio-medical, disposable pre-gelled silver chloride electrodes; GS-26) were placed around participant’s left forearm. When electrodes were poorly gelled, additional electrode conductant gel was applied. The first electrode pair was placed on the axis of the left extensor digitorum muscle; the remaining electrode pairs were placed around the forearm, roughly equidistant from each other with a preference for sites with optimal muscle contact. Additionally, a single reference electrode was placed on the elbow, above the olecranon bone. To hold the electrodes in place throughout the training, a sweatband was placed around the electrodes. Markings on the skin were used to ensure stable electrode positioning across sessions.

#### Calibration protocol

Bionic hand gestures were introduced to participants serially over the training sessions. On D1, participants calibrated the EMG controller on the bionic open and close gestures. On D2, participants added bionic pinch to their controller. On D3, participants added bionic tripod to their controller.

At the beginning of training day 1 (D1), participants practiced making natural muscle contractions for the first two hand gestures in each respective control strategy (biomimetic: open and close; arbitrary: 1 finger and 2 fingers). During calibration, participants were instructed to execute hand gestures at 20% of their maximum voluntary contraction (e.g. *“Would you be able to continue to use this level of force 1000 times in the session?”*). Using the Coapt COMPLETE CONTROL system [Gen2; (*18*)], participants calibrated their EMG controller by serially performing each gesture, guided by the experimenter. This auto-calibration process recorded EMG data during the muscle contractions for each gesture and auto-segmented and auto-labelled the EMG data for each bionic hand gesture class. To maximize the generalizability of the calibration data to the training tasks, we utilized Coapt’s Adaptive Advance feature (*47*, *48*) which implements a layering-like algorithm to combine multiple sets (layers) of training data for each gesture. Therefore, we added 7 layers of training data for each gesture: 3 layers with the arm positioned in front of the subject at a 90 degree angle, 1 layer with the arm positioned to the left, 1 layer with arm positioned to the right, 1 layer with the arm positioned upright and 1 layer with the arm positioned back at baseline. The Coapt system used established classification parameters including: 200 ms analysis windows with a 25 ms update increment, time domain and auto regressive features extracted from each window. Subsequent data were then classified by a linear discriminant analysis classifier (*49*, *50*). Bionic hand speed control was proportional to the EMG activity (*51*). However, the intention to move is constrained by the motors of the bionic hand, creating a delay.

### Training protocol

Between D1-D4, participants completed a series of tasks designed to quantify different aspects of bionic hand skill: speed, dexterity and gesture switching. For all tasks, participant’s training performance was filmed for an experimenter to perform an offline analysis of all relevant measures.

#### Speed (Southampton Hand Assessment Procedure)

At the beginning of every training session, participants completed a modified version of the SHAP (*52*). Participants were instructed to transfer an object as quickly as possible from one position to another. After each transfer, the experimenter would return the object to the starting position. On D1, participants performed 20 speed transfers of lightweight and heavyweight cylinder objects over 6cm using the open and close bionic hand gestures. On D2, participants repeated the same trials as in D1, and 20 transfers of lightweight and heavyweight ‘tip’ objects over 5cm using the open and pinch bionic hand gestures. On D3, participants repeated the same trials as D2, and then 20 transfers of the lightweight and heavyweight ‘tripod’ objects over 5cm using the bionic hand open and tripod gestures. On D4, participants repeated the same trials as in D3, and then 20 transfers of the lightweight ‘tripod’ objects using the bionic hand open and tripod gestures. On the generalization session, participants performed 20 speed transfers of the lightweight cylinder object using the bionic hand open and close gestures

One participant’s speed data was excluded, due to a technical issue with their Coapt EMG controller. This resulted in their control speed being over 3 standard deviations slower than the group average.

#### Dexterity (Virtual Eggs Test)

The Virtual Eggs Test is a modified version of the box and blocks test (*53*). It involves fragile blocks [i.e, “eggs”; (*54*, *55*)]. The virtual eggs (40×40×40mm, ~80g) exploit a magnetic fuse mechanism which collapses (i.e., breaks) when grasped with a grip force larger than a specific threshold. The break point was calibrated at a force value that was roughly 6N.

During the task, participants were instructed to transfer eggs over a 20cm tall wooden wall as fast as possible without breaking them. The task was only performed using the open and close bionic hand gestures. Participants were encouraged to prioritize grasping the eggs successfully over speed. Participants were told that if the egg broke on initial grasp, they were required to still complete the transfer. Performance was measured as the number of unbroken (successful) eggs transferred within a 2-minute time-period. Prior to starting the task, participants were allowed to practice transferring one egg. Additionally, using a customized glove for the bionic hand, flexiForce sensors [B201-M-8; Tekscan (*56*)] measured force applied on the thumb and index finger pads throughout the task. The sensor has a 0.375-inch sensing area diameter. Pressure data was recorded using the ELF System (Tekscan).

#### Gesture switching (block stacking)

Participants had to learn how to engage the gesture switching functionality of the bionic hand. To switch into a new gesture, participants would first need to engage an open hand signal. This would automatically trigger the bionic hand to move into a baseline hand open position, ready to switch. Participants could then perform a muscle contraction associated with the desired bionic gesture (close, pinch, tripod) they wanted to switch into. If a short sequence of the correct signal was sent, exceeding a gesture selection confidence threshold, the bionic hand would automatically switch into the open version of the desired bionic gesture and lock into that gesture until switched again. Any maintained signal of the grasping gesture would close the bionic hand proportionally into that closed version of that locked gesture.

To quantify the ability for users to successfully perform gesture switching, we designed a block stacking task that required participants to grab blocks using pre-defined bionic hand gestures. There were two variations of the task. The 2 gesture version required participants to switch between close and pinch. The 3 gesture version required participants to switch between close, pinch and tripod. Participants performed the 2 gesture version on D2, D3 and D4 and the 3-gesture version only on D3. Prior to starting the task, participants were instructed to grab, transfer and stack blocks (large blocks: 2×2×2in; small blocks: 1×1×1in) into towers of 3, as quickly as possible. There were 4-blocks for each of the gestures being tested (i.e., 8 total blocks for the 2-gesture version; 12 total blocks for the 3-gesture version). The blocks were arranged such that participants would have to grab a block with the first gesture (close) and the next block with the next gesture (pinch), and so on. If a participant was in an incorrect gesture, participants were instructed to try again until correct. The task finished when participants had successfully transferred all blocks.

### Pre-post testing protocol

We also used a set of pre-post comparison testing measures assessed before and after training: control automaticity, motor control, classification accuracy, and sense of embodiment.

#### Automaticity of bionic hand control

##### Cognitive load task

To assess the cognitive load imposed by bionic hand use, a concurrent numerical cognition task was performed during the first (D1) and last (D4) training sessions. The task was adapted from previous studies (*34*). Participants were asked to perform two variations of a block stacking task. The single condition task required participants to quickly grab, transfer, and stack as many blocks (2×2×2in) as possible into towers of three using the bionic hand. The dual condition task required participants to perform the same block stacking task, while simultaneously verbally performing a counting task. The counting task required participants to follow along to a set of low, medium, and high pitch auditory tones played from a laptop. The tones were presented every 2 to 4 s in a randomized order, for a total duration of 1 min. Participants started the task with the initial count of “50”. Participants were then instructed to: (i) add 3 to the current number if they heard a high pitch tone, (ii) hold the current count if they heard a medium pitch tone, or (iii) subtract 3 to the current count if they heard a low pitch tone. For each sound, participants were instructed to verbally respond. To ensure participants were equally motivated for their motor and counting performance, participants were told that their performance would be scored equally on the number of blocks transferred and their counting performance. The primary measures we analyzed were the number of blocks participants transferred in the single and dual condition tasks (separately), and their counting accuracy (i.e., how many trial counts were correct).

To obtain a baseline for counting performance irrespective of motor performance, participants first performed the counting task without any block stacking. Next, participants performed the dual condition task. Finally, to obtain a baseline for motor performance without cognitive load, participants performed the single condition (block stacking only) task. In this latter condition, the counting sounds were still played throughout the task however, participants were told to ignore them.

For each participant, we first calculated the total number of blocks transferred in the single and dual condition tasks (separately). To quantify how cognitively demanding the numerical cognition task was on bionic hand motor performance, a *Control Automaticity* ratio was computed by dividing the number of blocks transferred in the dual condition task by the number of blocks transferred in the single condition task. Counting performance was computed by taking the percentage of total correct mathematical operations.

##### Control difficulty questionnaire

At the end of every session, participants were asked to respond to the following question: *“How difficult was it to control the prosthesis? Please rate between 0 (I found it as easy to perform the movement as using my own hand) to 10 (the most difficult thing imaginable).”*

#### Motor control

To quantify participants’ motor control for both their left biological hand and the bionic hand, participants performed a ballistic reaching task during the familiarization session and the last (D4) training session (see Supp. Figure 1). Untrained participants completed this task during the familiarization session and the beginning of the generalization session.

Participants were seated at a custom-made wooden tabletop placed above a digitizing tablet (42.6 by 28.4 cm, Intuos Pro Large; Wacom, Vancouver, WA) and facing an LCD monitor (15.6in, 1920 x 1080 pixel dimension; Dell Precision 3560). The participants performed reaching movements by sliding a digitizing stylus (Wacom Pro Pen 3D; Wacom, Vancouver, WA) across the tablet. The position of the stylus was recorded by the tablet at 60Hz. The experimental software was custom written in python for PsychoPy (v2021.1.1). Direct vision of the arms (elbows and shoulders included) was occluded using a black barber cape. Additionally, the lights were extinguished in the room to minimize peripheral vision of the hand.

Participants performed center-out planar reaching movements to visual targets. Due to the time constraints with fitting and removing the bionic hand system, all participants performed the task first with their biological hand and then with the bionic hand. Prior to starting the task, the experimenter locked the bionic hand around the digitizing stylus so that it was immoveable (could not open) for the task. Participants were shown their hand position, the home location and, at during trials, the reach target. The hand position was constantly shown to participants (60Hz), indicated by a green crosshair (0.36cm x 0.36 cm). The home location was constantly shown to participants as a square (0.36cm x 0.36cm) at the bottom, center of the screen (1.8cm above the bottom of the screen). The home location was colored grey between trials and red during trials. The reach target was a white circle (0.18cm radius) that would appear at 3 separate locations (left, center, right). The left and right targets were 67 degrees from vertical center. All targets were 12.8cm away from the home location. We also displayed the trial number in the top right corner of the monitor.

Participants completed 10 practice trials, prior to starting the task. Participants completed 60 experimental trials. A trial was initiated once participants hovered the cursor over the home location. The home location would then turn red to denote that a reach targe would soon appear in either one of three target locations. After 3 seconds, a reach target would appear in one of the 3 locations. Participants were instructed to perform a fast, ballistic reaching movement (within 2 seconds) towards the center of each target and to avoid corrective movements, such that they should maintain the end position of their reach until the target disappeared. In total, each trial lasted 5 seconds. Participants were then required to return to the home location to begin the next trial. The three reach targets were each presented 20 times. The order of the targets was pseudorandomized, such that each target was randomly sampled in batches of 3. All participants were presented with the same trials order.

Due to technical issues, we excluded the first experimental trial for all participants (i.e., 59 total experimental trials). Additionally, trials were excluded where the reach end point was above or below 2 standard deviations of the subject’s reach error (range of excluded trials across subjects was 1-6% of total trials).

#### Classification accuracy

To quantify participants’ classification accuracy, we used the real-time signal classification output from the Coapt EMG controller [Gen2; (*18*)]. Participants were seated at a table, facing an LCD monitor (15.6in, 1920 x 1080 pixel dimension; Dell Precision 3560). Active control of the bionic hand was turned off. While the bionic hand could not move, the EMG controller was still turned on. On the monitor, participants were shown their real-time signal classification (frame rate: 50ms), listed as words: “Open”, “Close”, “Pinch”, “Tripod” or no output, indicating “Rest”.

Participants were instructed to perform and maintain 6 different hand gestures each for 20 seconds. On the monitor, participants were given a visual indication of the start and end of each gesture trial, as well a virtual expanding circle proportional to their level of contraction (Coapt virtual game: In the Zone). Participants were instructed to perform and maintain each gesture such that they could maintain correct classification, on the monitor, for each of the desired gestures for the duration of the trial. When assessing classification accuracy for 3 gesture classes, the trial order was rest, open, close, rest, open close. When assessing classification accuracy for 5 gesture classes, the trial order was rest, open, close, rest, pinch, tripod. Note that when running this task, open bionic gesture trials would terminate after 2.5 seconds if correct classification and a specified force level was maintained (due to unrelated purposes for a separate study).

The task output file included (i) the real-time classification decision, (ii) the trial number the participants were on and (iii) the EMG activity for each of the 8 channels (all updated every 50ms). Previous research has taken a variety of approaches to define a relevant time window for offline analysis of classification accuracy (*57*). Because “Open” trials would sometimes terminate early (as noted in the previous paragraph), we employed an analysis approach that could control for differences in overall trial duration between gestures. We opted to use the first correct classification point, for each gesture trial, as timepoint 0. Therefore, the average classification accuracy was computed from timepoint 0 to 49 (approximately the first 2.5 seconds of each 20 second gesture). This analysis approach was performed for each gesture separately for each participant. On the 3-gesture version of the task, where participants were asked to perform each gesture once (i.e., 2 trials per gesture), classification accuracy was calculated separately for each trial of the same gesture and then those values were averaged. To construct the group-level confusion matrices in Supp. Figure 2, values were then averaged across participants for each group. To compute a single value of average classification (shown in Supp. Figure 2D,F), we took the average correct classification across gestures (i.e. diagonal of confusion matrix).

Due to technical issues at the beginning of the study, we were not able to acquire these data from the first 4 study participants (3 arbitrary, 1 biomimetic).

#### Sense of embodiment questionnaire

To assess changes in the sense of embodiment over the bionic hand, participants were asked to complete an embodiment questionnaire before the first and after the last training session (Supp. Table 1). Untrained participants filled the post-questionnaire out after completing the post-motor control assessments, prior to being able to experience active control of the bionic hand. The questionnaire was focused on the explicit (phenomenological) aspect of embodiment, whether the bionic hand feels like a part of one’s body. Participants were asked to rate their agreement with 10-statements (*34*, *58*) on a seven-point Likert-type scale ranging from −3 (strongly disagree) to +3 (strongly agree). Statements were clustered into three main categories, probing different aspects of embodiment: body ownership, agency, and body image. For each participant, questionnaire scores were averaged within each embodiment category. To compute a difference score, pre-scores for each embodiment category were subtracted from the post-scores.

### Statistical analysis

All statistical analyses were performed using JASP (v0.14). Tests for normality were conducted using a Shapiro-Wilks test. When assumptions of normality were met, we used parametric statistics, and when they were not met (p<0.05 for the Shapiro-Wilks tests), equivalent nonparametric tests were used. Between-group comparisons were conducted using repeated measure ANOVAs with group (biomimetic, arbitrary or untrained) as a fixed effect and independent samples t-tests (parametric) or Mann-Whitney tests (non-parametric). Within-group comparisons were conducted using paired t-tests (parametric) or Wilcoxon tests (non-parametric). All non-significant results were further examined using corresponding Bayesian tests under continuous prior distribution (Cauchy prior width r = 0.707). We interpreted the test based on the well accepted criterion of Bayes factor smaller than 1/3 (*59*) as supporting the null hypothesis.

## Acknowledgements

We thank John Ingelholm for technical support. Marco Controzzi and Francesco Clemente for sharing the original design of the virtual eggs. Lindsey Emmrich (Ossur) for consultation on the bionic hand system design and training paradigm. Anthony Moore (Ossur) and Nathan Wagner (Ossur) for fabrication of the bionic hand system. Lina Teichmann for providing technical and conceptual support throughout the study. The OP4 clinic nursing staff for their patience and support for our 60 participants. Mike Reel, our nurse practitioner, for his commitment and patience with all of the participant visits. The Medical Centre for Orthotics and Prosthetics (MCOP) for help in device fabrication and technical support. The technical support team at Coapt for providing regular technical support for the EMG controllers.

## Author contributions

H.R.S. designed the research, collected the data, analyzed all datasets and wrote the first version of the manuscript. T.R.M. and C.I.B. designed the research, supervised analyses and edited the manuscript. M.U. assisted in data collection, analyzed the motor control data and edited the manuscript. M.M. assisted in data collection, analyzed the classification accuracy data and edited the manuscript. B.R. assisted in data collection and edited the manuscript. J.V., B.L., and L.H. provided technical support and edited the manuscript.

## Data and materials availability

Pre-registered study predictions and methods, as well as the data used in the study, can be accessed at: https://osf.io/3m592/. Code used in the study can be accessed at https://github.com/hunterschone/ProControl.

